# Intrinsically disordered regions of Tristetraprolin and DCP2 directly interact to mediate decay of ARE-mRNA

**DOI:** 10.1101/2021.11.07.467627

**Authors:** Vincent D. Maciej, Nevena Mateva, Theresa Dittmers, Sutapa Chakrabarti

**Author notes:** To whom correspondence should be addressed: Sutapa Chakrabarti.

## Abstract

The RNA binding protein Tristetraprolin (TTP) is a potent activator of mRNA decay, specifically for transcripts bearing AU-rich elements (AREs) in their 3′-untranslated regions. TTP functions as a mediator for mRNA decay by interacting with the decay machinery and recruiting it to the target ARE-mRNA. In this study, we report a weak, but direct interaction between TTP and the human decapping enzyme DCP2, which impacts the stability of ARE-transcripts. The TTP-DCP2 interaction is unusual as it involves intrinsically disordered regions (IDRs) of both binding partners. We show that the IDR of DCP2 has a propensity for oligomerization and liquid-liquid phase separation (LLPS) *in vitro*. Binding of TTP to DCP2 leads to its partitioning into phase-separated droplets formed by DCP2, suggesting that molecular crowding might facilitate the weak interaction between the two proteins and enable assembly of a decapping-competent mRNA-protein complex on TTP-bound transcripts in cells. Our studies underline the role of weak interactions in the cellular interaction network and their contribution towards cellular functionality.

## Introduction

Degradation of messenger RNA (mRNA) serves as an important mechanism for achieving rapid changes in the gene expression profile of a cell in response to external stimuli. Half-lives of mRNA can vary drastically in eukaryotes: transcripts encoding cytokines, proto-oncogenes and transcription factors are highly labile and are degraded within minutes, while those of house-keeping genes are stable over hours (Khabar 2005). The intrinsic stability of an mRNA transcript is often determined by specific sequences or structural elements that are usually located in the 3′-untranslated region (3′-UTR) of the mRNA. These *cis*-acting elements are recognized by distinct *trans*-acting protein factors, which in turn recruit the mRNA degradation machinery to the transcript to ensure its timely decay (Mayr 2017). A predominant *cis*-acting element in higher eukaryotes is the AU-rich element (ARE), a nonameric sequence motif rich in adenines and uridines found in multiple copies in the 3′-UTR, which is recognized with high specificity and affinity by AU-binding proteins (AUBPs) (Khabar 2017).

Of the AUBPs competent in triggering mRNA decay, the most extensively studied are the protein Tristetraprolin (abbreviated as TTP and also known as Tis11/ZFP36) and its paralogues BRF1 (Tis11b/ZFP36L1) and BRF2 (Tis11d/ZFP36L2) (Wells, Perera et al. 2017). TTP and its paralogues are characterized by the presence of two CCCH zinc finger motifs arranged into a tandem zinc finger (TZF) domain, which is flanked at the N- and C-termini by long stretches of low-complexity sequences (Figures 1A and S2D). The central TZF domain binds RNA with high affinity and is fairly well-conserved among the paralogues (Hudson, Martinez-Yamout et al. 2004), while the flanking regions diverge considerably in length and sequence (Sequence alignment, S1A). The N- and C-terminal stretches are referred to as “activation domains” for their role in recruiting components of the mRNA degradation machinery to activate mRNA decay. Co-immunoprecipitation studies indicate that TTP and BRF1 are capable of recruiting the CCR4-NOT deadenylase complex and components of the RNA exosome, as well as the decapping machinery (DCP1, DCP2 and Edc3) and the 5′-3′ exonuclease, Xrn1 (Chen, Gherzi et al. 2001, Fenger-Gron, Fillman et al. 2005, Lykke-Andersen and Wagner 2005, Hau, Walsh et al. 2007). The N-terminal activation domain of TTP was shown to be important for decapping of an ARE-substrate, while the C-terminal domain engages the CNOT1 protein to recruit the CCR4-NOT complex (Fenger-Gron, Fillman et al. 2005, Sandler, Kreth et al. 2011, Fabian, Frank et al. 2013). Recruitment of the CCR4-NOT complex to an RNA substrate establishes a vast interaction network, involving factors that mediate decapping and translational repression, in addition to deadenylation and subsequent 3′-5′ exonucleolytic decay (Nishimura, Padamsi et al. 2015). Therefore, a direct interaction of TTP with CNOT1 is an efficient means of triggering rapid mRNA decay. The NOT1-binding region of TTP was mapped to a conserved sequence motif at its very C-terminus (denoted as NIM in Figures 1A and S1A) (Fabian, Frank et al. 2013). Interestingly, a TTP construct lacking this motif was still capable of mediating decay of an ARE-substrate, suggesting that additional motifs in TTP engage with the degradation machinery independently of CNOT1 to facilitate mRNA decay, and ensure redundancy in this process (Sandler, Kreth et al. 2011, Fabian, Frank et al. 2013).

**Figure 1.**
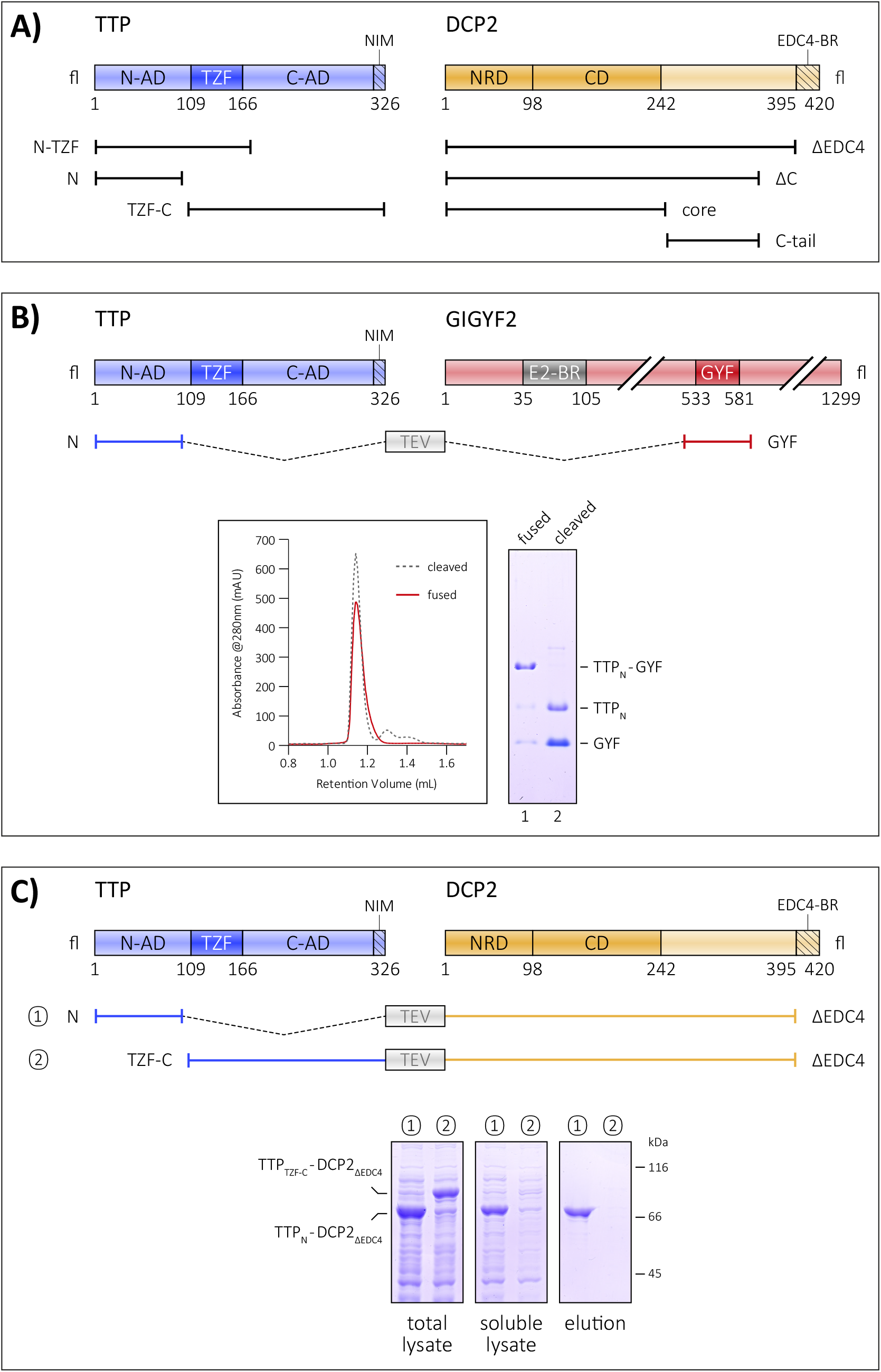
Interaction of the TTP N-terminal activation domain with GIGYF2 and DCP2. A) Schematic representation of the domain organization of TTP and DCP2 and the constructs used in this study. The central tandem zinc finger domain of TTP (dark blue) is flanked by the unstructured N- and C-terminal activation domains (N-AD and C-AD, respectively) on either side (light blue). The structured core of DCP2 (dark orange) comprises its N-terminal regulatory domain (NRD) and its catalytic Nudix domain (CD) and is followed by a long, unstructured C-terminal region (light orange), the last 25 residues of which harbor a binding motif for EDC4 (EDC4-BR, indicated by dashed lines). B) The GYF domain of GIGYF2 restores proteolytic stability of the TTP N-terminal activation domain (TTP_N_). A schematic representation of GIGYF2 indicates the position of the GYF domain (dark red) in the primary structure of the protein, with respect to other binding motifs, such as the eIF4E2-binding motif (in grey, denoted by E2-BR). The fusion polypeptide of TTP_N_ and GYF, separated by a linker sequence bearing a TEV protease cleavage site is shown. Analytical size-exclusion chromatography (SEC) and the corresponding SDS-PAGE analysis of peak fractions depict the stability of TTP_N_ in complex with GYF. The retention volume of the single polypeptide (fused) is identical to that of non-covalently linked complex (cleaved, referring to cleavage by TEV protease), indicating formation of a stable complex. TTP_N_ in complex with GYF is stable even upon TEV-cleavage of the fusion polypeptide. C) Schematic representation of the TTP-DCP2 fusion polypeptides, designed similar to the TTP_N_-GYF fusion described above. Comparison of the stability of TTP_N_-DCP2_ΔEDC4_ with that of TTP_TZF-C_-DCP2_ΔEDC4_ by Ni^2+^-affinity pulldowns. Both fusion proteins are well-expressed but only TTP_N_-DCP2_ΔEDC4_ can be isolated by Ni^2+^-affinity purification, indicating that fusion of DCP2 rescues the behaviour of TTP_N_ but not TTP_TZF-C_. The left, middle and right panels indicate total lysate, soluble lysate and elution, respectively. Total lysate and soluble lysate refer to samples collected before and after centrifugation of the cell-suspension after sonication.

To understand the contribution of the N-terminal activation domain towards TTP-mediated decay, we set out to investigate the interaction between TTP and the decapping enzyme DCP2. DCP2 is a bilobed protein consisting of an N-terminal regulatory domain (NRD) and a catalytic Nudix domain (CD) that mediates hydrolysis of the 7-methyl guanosine cap of eukaryotic mRNA (She, Decker et al. 2008, Mugridge, Ziemniak et al. 2016, Valkov, Muthukumar et al. 2016, Wurm, Holdermann et al. 2017). The structured core is followed by an unstructured C-terminal extension that harbors motifs for anchoring proteins such as the enhancer of decapping 4 (EDC4, also known as Hedls), which is a scaffold for assembly of decapping factors in metazoans (Figure 1A) (Chang, Bercovich et al. 2014). We found a weak, but direct interaction between the TTP N-terminal activation domain and DCP2 that is mediated by intrinsically disordered regions (IDRs) of both proteins and show that decay-activation by the TTP N-terminal domain is dependent on DCP2. We propose a mechanism for how molecular crowding in cells might promote the weak interaction between TTP and DCP2 and facilitate assembly of decay-competent messenger ribonucleoprotein complexes (mRNPs) on ARE-transcripts.

## Results

### The N-terminal activation domain of TTP associates with the decapping enzyme DCP2

*In vitro* biochemical and biophysical studies on human TTP have largely been marred by the proteolytically unstable nature of the full-length protein as well as shorter constructs spanning the N- and C-terminal activation domains. Previous structural and biochemical studies have relied on short synthetic peptides corresponding to binding motifs and fusions with the maltose-binding protein (MBP) to stabilize the protein, although other commonly used stability tags such as glutathione-S-transferase (GST), GFP and thioredoxin do not have a similar effect (Figure S1B). We hypothesized that binding of an interaction partner to TTP would minimize its proteolytic degradation and stabilize the protein. The translational repressor GIGYF2 is a known interactor of TTP, with its GYF domain directly binding the TTP-PPPG_ϕ_ motifs (Fu, Olsen et al. 2016). Therefore, we designed a construct where the N-terminal activation domain of TTP (TTP_N_) containing a PPPGF motif was fused to the GIGYF2-GYF domain (hereafter referred to as GYF) to generate a single polypeptide, with the two interacting partners connected by a short linker bearing a 3C protease-cleavage site (Figure 1B). The resultant fusion construct is remarkably stable and yields large amounts of a homogenous species. Interestingly, TTP_N_ remains stable even after cleavage of the polypeptide with 3C protease to yield a non-covalently associated TTP-GYF complex, indicating that most of the proteolytic degradation of TTP occurs during its expression (Figure 1B, lane 2). Encouraged by this observation, we proceeded to generate fusion constructs of the TTP N- and C-terminal activation domains with DCP2 (Figure 1C). As full-length DCP2 is also proteolytically unstable, a C-terminal truncation of DCP2 lacking the 26-residue EDC4 binding-region (DCP2_ΔEDC4_) was used instead (Figure 1A). Fusion of DCP2_ΔEDC4_, akin to GYF, conferred stability on TTP_N_ and led to the isolation of highly pure fusion protein, but did not have a stabilizing effect on the TTP C-terminal activation domain (Figure 1C). The TTP_C-TZF_-DCP2_ΔEDC4_ fusion protein is expressed in an amount comparable to TTP_N_-DCP2_ΔEDC4_, but little to no protein was observed in the soluble lysate and the elution of the small-scale Ni^2+^-affinity pulldown, suggesting that the protein is unstable in solution. Our results suggest a direct interaction between TTP_N_ and DCP2, consistent with previous reports of the N-terminal activation domain of TTP engaging the decapping machinery.

### The IDRs of TTP and DCP2 mediate a direct but weak interaction between the proteins

In order to determine the region or domain of DCP2 that mediates stabilization of TTP, we designed a series of constructs where TTP_N_ is linked to different DCP2 constructs (Figure 2A). As mentioned above, fusion of TTP_N_ to DCP2_ΔEDC4_ results in a stable protein (Figures 1C and 2B). Truncation of an additional 40 residues from the C-terminus of the DCP2_ΔEDC4_ construct (DCP2_ΔC_) does not mitigate its ability to stabilize TTP (Figure 2B, compare lanes 1 and 2). The DCP2 NRD-Nudix core does not stabilize TTP_N_ to the same extent as the other constructs, as the amount of TTP_N_-DCP2_core_ protein obtained in the soluble lysate and elution of the Ni^2+^-affinity pulldown is significantly lower than for the other fusion constructs (Figure 2B, compare lane 3 to lanes 1, 2 and 4). Moreover, the TTP_N_-DCP2_core_ appears to degrade during subsequent purification steps (Figure S2A), unlike fusions bearing the DCP2 C-tail which remain stable over time. Taken together, our observations indicate that the binding motif for TTP resides in the C-terminal extension of DCP2 that is proximal to its NRD-Nudix core (referred to as C-tail). Consistently, fusion of the DCP2 C-tail is sufficient to confer stability on TTP_N_ (Figure 2B, lane 4).

**Figure 2.**
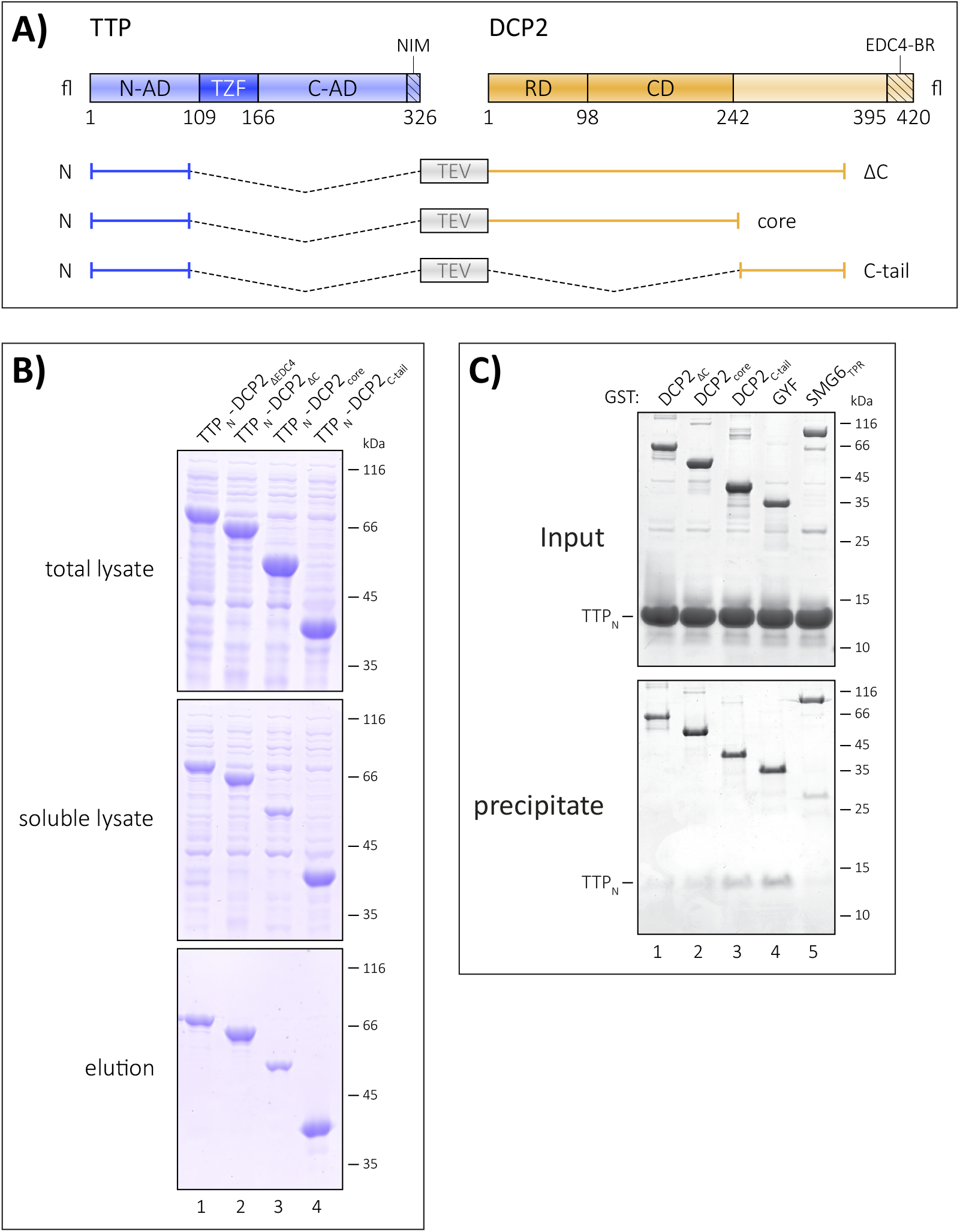
The TTP-binding site resides in the C-terminal intrinsically disordered region of DCP2. A) Schematic representation of the fusion constructs comprising TTP_N_ and the different regions of DCP2. As in Figure 1, all fusion constructs contain a linker with a TEV protease cleavage site separating the TTP and DCP2 sequences. B) Ni^2+^-affinity pulldowns to compare the effect of the DCP2 NRD-CD core and the unstructured C-terminus on the stability of TTP_N_. The top, middle and bottom panels indicate total lysate, soluble lysate and elution, respectively. Fusion of DCP2_core_ to TTP_N_ has the least impact on its proteolytic stability (see also Figure S2A), while all DCP2 constructs bearing the C-tail have a remarkable stabilizing effect. C) GST-pulldown assays of isolated TTP_N_ with GST-DCP2 proteins. GST-GYF and GST-SMG6_TPR_ serve as positive and negative controls, respectively. The top and the bottom panels indicate input and precipitate, respectively. In this experiment proteins have been visualized by silver staining. GST-DCP2_C-tail_ co-precipitates TTP_N_ to a similar extent as GST-GYF, while the amount of TTP_N_ co-precipitated with GST-DCP2_core_ is negligible. Despite containing the C-tail, GST-DCP2_ΔC_ fails to co-precipitate TTP_N_, possibly due to folding back of the C-tail on the NRD-CD core (see also Figure S2B).

To further investigate the association of TTP and DCP2, we performed GST-pulldowns using GST-tagged DCP2 proteins and untagged TTP_N_ (Figure 2C). The NMD endonuclease SMG6_TPR_ and GYF were used as negative and positive controls for this assay, respectively. As anticipated, TTP_N_ co-precipitated with GST-DCP2_C-tail_, but did not bind GST-DCP2_core_ as much (Figure 2C, compare lanes 3 and 2). Surprisingly, the DCP2_ΔC_ construct, which was capable of stabilizing TTP upon fusion, did not co-precipitate any TTP_N_ protein. We attribute this to an interaction between the DCP2 core and its C-tail, which in an intramolecular context (in DCP2_ΔC_) takes precedence over the intermolecular TTP_N_-DCP2_C-tail_ interaction (Figure S2B). Indeed, the inability of DCP2_ΔC_ to mediate stable interactions with TTP_N_ in *trans* allowed us to isolate large amounts of homogenous TTP_N_ from the parent fusion protein for our biochemical and biophysical studies (Figure S2C and Supplemental Methods). The interaction of TTP_N_ with DCP2_C-tail_ presents an unusual mode of interaction, where the binding regions of both interacting partners are predicted to be disordered (Figure S2D).

We next performed analytical size-exclusion chromatography assays to compare the interaction of TTP_N_ with GYF and DCP2_C-tail_. Incubation of TTP_N_ and GYF in a 1:1.2 molar ratio for 16 hours at 4°C resulted in formation of a stable complex, as indicated by a shift in peak retention volume in comparison to the individual proteins (Figure 3A, left panel). In contrast, TTP_N_ did not form a stable complex with DCP2_C-tail_, even at high concentrations of 100 μM, suggesting a weaker interaction between the two proteins (Figure 3A, right panel). The molecular weight of the TTP_N_-DCP2_C-tail_ fusion, as determined by size-exclusion chromatography coupled with multi-angle light scattering (SEC-MALS), is approximately about 26 kDa, and is reduced to approximately 11 kDa upon cleavage with TEV protease (Figure 3B). This indicates that unlike GYF, DCP2_C-tail_ does not remain associated with TTP_N_ upon cleavage of the fusion polypeptide, which is consistent with a weak interaction between the two proteins.

**Figure 3.**
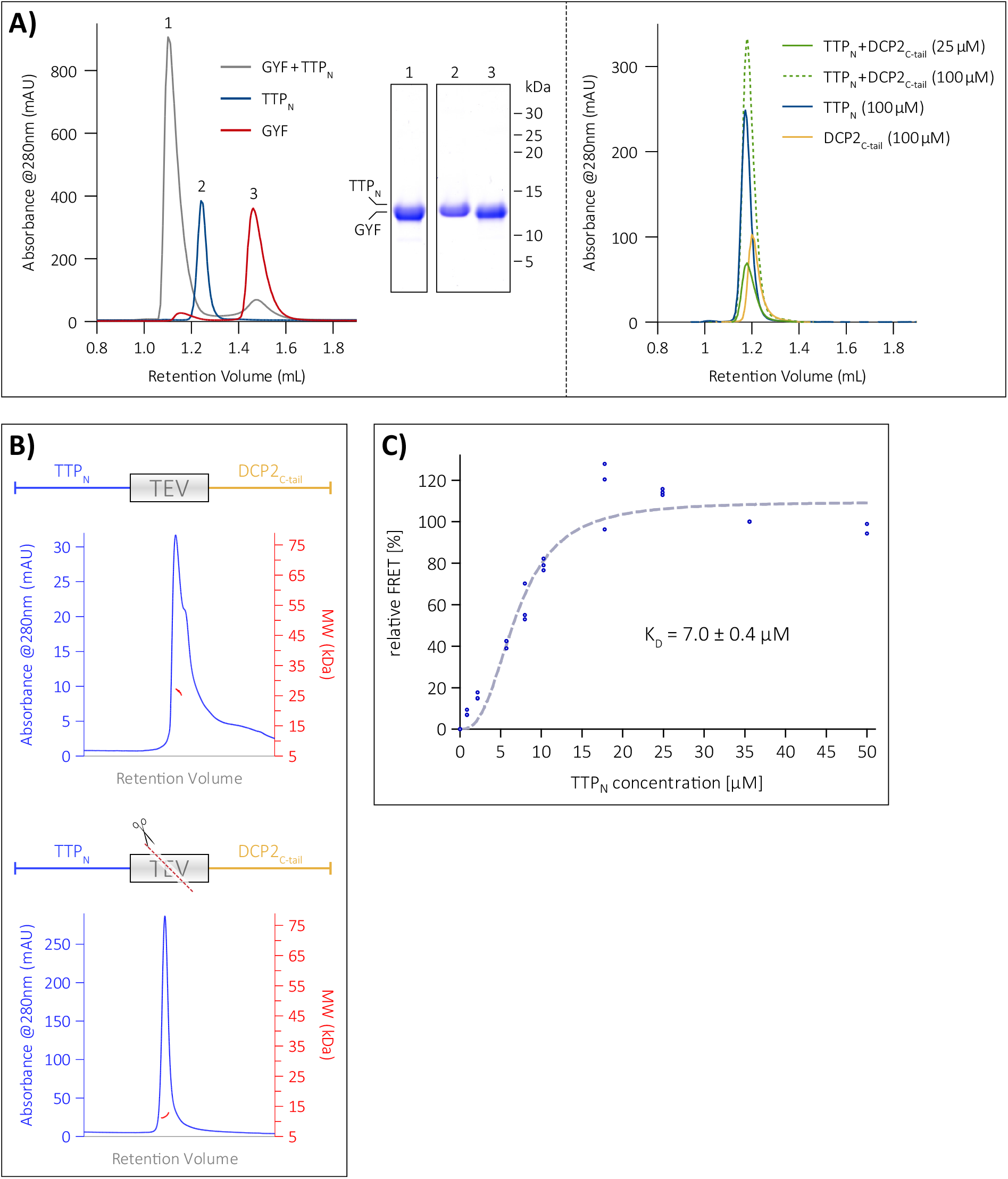
The N-terminal activation domain of TTP engages DCP2 in a low-affinity interaction. A) Analytical SEC analyses of equimolar mixtures of TTP_N_ and GYF (left panel), and TTP_N_ and DCP2_C-tail_ (right panel). SEC runs of the individual proteins (TTP_N_, DCP2_C-tail_ and GYF) are shown for comparison. TTP and DCP2 do not form a stable complex, even at reasonably high concentrations of 100 μM, indicating weak binding while the interaction of TTP with GYF is considerably stronger and results in a complex (see corresponding SDS-PAGE analysis of the peak fractions). B) Multi-angle light scattering in tandem with SEC (SEC-MALS) to analyze the TTP_N_-DCP2_C-tail_ fusion polypeptide in its intact (top) and TEV protease-cleaved (bottom) states. The intact fusion has a molecular weight of 26.5 kDa (in good agreement with the theoretical molecular weight of 23.2 kDa). Cleavage of the fusion by TEV protease results in a molecular weight of 11.6 kDa, which does not match that of the complex, but approximates the molecular weights of isolated TTP_N_ (11.1 kDa) and DCP2_C-tail_ (12.1 kDa). C) Intermolecular FRET assay using Cy3-DCP2_C-tail_ (donor) and Cy5-TTP_N_ (acceptor) to determine the equilibrium dissociation constant (*K*_D_) of the interaction. The mean relative FRET percentage calculated from 3 independent experiments (only 2 measurements for the 50 μM data point) were fitted to a one-site specific binding curve with Hill-slope. The resultant *K*_D_ of 7 μM indicates a low-affinity interaction. All data points are shown in the figure. Plotting of the data points and curve-fitting were performed using Graphpad Prism 5.0.

To quantify the strength of the interaction between TTP and DCP2 in solution, we used Förster Resonance Energy Transfer (FRET), which allows measurements over a wide range of binding affinities. DCP2_C-tail_ and TTP_N_ were labelled at a single position with the fluorescent dyes Cy3 and Cy5, respectively (Figure S3A). An increasing amount of the acceptor fluorophore, Cy5-TTP_N_, was titrated into a constant amount of the Cy3-DCP2_C-tail_ donor, with the unlabeled TTP and DCP2 proteins serving as controls in the FRET setup. The tendency of DCP2_C-tail_ to oligomerize at high concentrations under non-reducing conditions, as observed in SEC-MALS (Figure S3B), precluded its use as an acceptor at high concentrations in FRET. Saturation of relative FRET efficiency with increasing concentrations of TTP_N_ suggests a specific binding to DCP2_C-tail_ with an estimated dissociation constant (*K*_D_) of 7 μM (Figure 3C).

### Functional impact of the N-terminal domain of TTP

The modest affinity of the interaction between TTP and DCP2 necessitates that these proteins are present in sufficiently high amounts in cells at a given point of time to ensure binding when the need arises. However, a quantitative study of protein abundance in HEK293 cells suggests that DCP2 is present in relatively low amounts, in comparison to other decapping and decay factors such as DCP1, EDC3, EDC4 and CNOT1 (Cho, Cheveralls et al. 2021). TTP and its paralogues are highly unstable and present in low amounts in uninduced cells. A possible mode of circumventing the low cellular concentration of a protein is to increase its local concentration by sequestering it into a cellular compartment. Two cytoplasmic compartments associated with mRNA processing (decay, storage or translational repression) are processing bodies (P-bodies) and stress granules (Decker and Parker 2012, Aizer, Kalo et al. 2014). These membrane-less compartments are assembled through liquid-liquid phase-separation, usually induced by multivalent, low-affinity protein-protein/RNA interactions (Nott, Petsalaki et al. 2015, Wang, Choi et al. 2018). These compartments are dynamic in nature and enable rapid recruitment and release of their components. DCP2 is a core component of P-bodies and is also present in stress granules (Sheth and Parker 2003, Teixeira and Parker 2007, Hubstenberger, Courel et al. 2017, Youn, Dunham et al. 2018, Youn, Dyakov et al. 2019). Interestingly, the N- and C-terminal activation domains of TTP were also shown to be capable of localizing to P-bodies, while TTP was found to be a component of stress granules (Franks and Lykke-Andersen 2007). We set out to investigate if the weak intra- and inter-molecular interactions mediated by the IDRs of DCP2 and TTP have an impact on their localization to membrane-less compartments. To this end, we first analyzed the tendency of DCP2_ΔC_, DCP2_core_, DCP2_C-tail_ and TTP_N_ to phase-separate into liquid droplets *in vitro* by phase-contrast microscopy. Consistent with IDRs playing a role in phase-separation and the tendency of the DCP2_C-tail_ protein to form higher order oligomers, we observed that DCP2_C-tail_ formed droplets upon addition of a crowding reagent (PEG8000). The droplets showed typical characteristics of liquid-liquid phase separation, such as growth, merging and reversible deformation. The DCP2 NRD-CD core also showed phase-separation under these conditions, although it does not contain long stretches of IDRs. Not surprisingly, DCP2_ΔC_, which contains both the core and the C-tail, displayed the strongest propensity to form droplets, even at a relatively low protein concentration (Figure 4A, top three panels). However, TTP_N_, which is predicted to be disordered, did not undergo phase-separation *in vitro* (Figure 4A, bottom panel). We next tested if the weak interaction between TTP_N_ and DCP2_C-tail_ in solution leads to recruitment of TTP to phase-separated droplets. In order to track both proteins, we labelled TTP_N_ and the DCP2 proteins with the fluorophores Cy3 and XFD488, respectively, and mixed them in equimolar ratios. As expected, all DCP2 proteins formed phase-separated droplets (Figure 4B). Interestingly, TTP_N_ partitioned into phase-separated droplets formed by DCP2_ΔC_ and DCP2_C-tail_ (as indicated by an overlap between the Cy3 and XFD488 signals), but not into those formed by DCP2_core_ (Figure 4B). Our observations suggest that the previously reported localization of the TTP N-terminal activation domain to P-bodies in cells might be a consequence of its weak interaction with DCP2.

**Figure 4.**
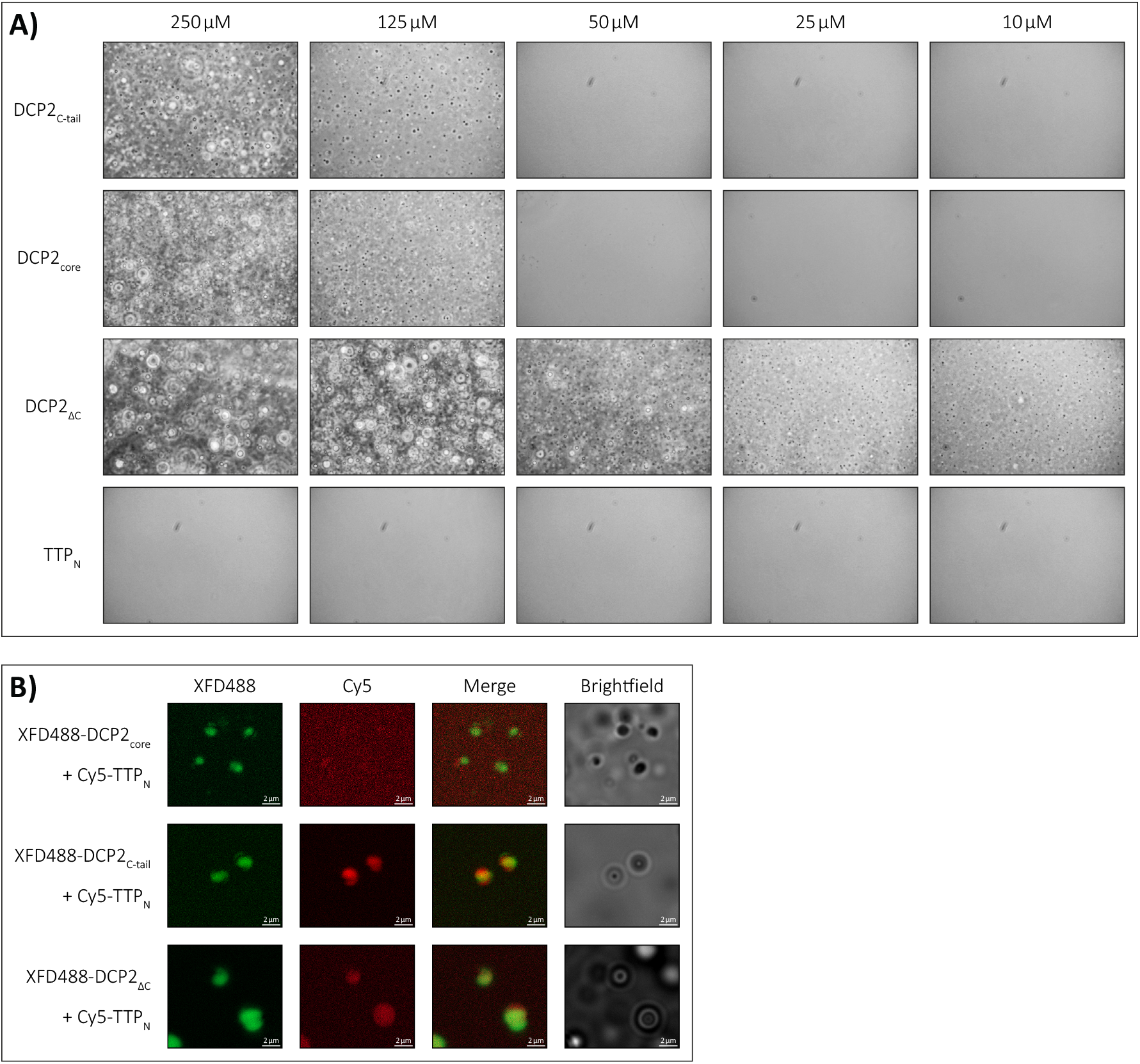
The low-affinity interaction of TTP with DCP2 induces LLPS of TTP_N_ in vitro. A) Representative images of *in vitro* liquid-liquid phase separation (LLPS) of the DCP2 proteins and TTP_N_ in presence of a molecular crowding reagent, PEG8000. A range of protein concentrations from 250 μM to 10 μM were analysed. All DCP2 proteins undergo LLPS at high concentrations, while DCP2_ΔC_ which has the strongest tendency to undergo phase separation, also does so at the lowest concentration investigated. TTP_N_ does not form phase-separated droplets in any of the tested conditions. Images were acquired by phase-contrast microscopy. B) Representative images of *in vitro* phase separation of equimolar mixtures of Cy5-labelled TTP_N_ and XFD488-labelled DCP2. Overlap of Cy5 and XFD488 signals suggests co-localization of the two proteins. TTP_N_ partitions into phase-separated droplets in the presence of DCP2_C-tail_ and DCP2_ΔC_, but not in the presence of DCP2_core_. Brightfield images show additional phase-separated droplets in different focal planes that are not visible in the fluorescent images due to optical sectioning by the Zeiss Apotome. The imperfect overlap between the XFD488- and Cy5-fluorescent images stems from moving droplets and a time-lag in image acquisition. Scale bars, 2 μm.

The observed low affinity interaction between TTP_N_ and the DCP2_C-tail_ also raises the question of whether the N-terminal activation domain of TTP plays a role in ARE-mediated decay, independent of the C-terminal NIM region. To assess this, we used a Renilla-luciferase (R-Luc) reporter containing two copies of the ARE motif of TNFα in its 3′-UTR, followed by the 3′-end of the non-coding RNA MALAT1 (Peter, Weber et al. 2017). The levels of R-Luc activity in HEK293 cells in the presence of different TTP constructs were measured as a readout of mRNA stability. A Firefly-luciferase (F-Luc) reporter was used as a transfection control, while an R-Luc reporter lacking the ARE motifs was used as a control for ARE-specific decay (Figure 5A). Comparison of R-Luc activity showed that the stability of the R-Luc-ARE reporter in the presence of TTP_N-TZF_ is approximately 1.5-fold higher than in the presence of full-length TTP, but considerably lower than in the presence of GFP, which served as a negative control (Figure 5A). This suggests that the TTP N-terminal activation domain is capable of mediating decay of ARE-RNA, independent of the C-terminal activation domain and the NIM-sequence therein. A TTP construct lacking the N-terminal activation domain was not tested in these assays as the nuclear export signal of TTP resides within its N-terminus (Phillips, Ramos et al. 2002, Lai, Wells et al. 2019), and its absence would render the protein incapable of localizing to the cytoplasm. Since TTP_N_ interacts with the DCP2_C-tail_, we next proceeded to test if the ability of the TTP N-terminal domain to activate decay is dependent on DCP2. To this end, we transfected HEK293 cells with a small interfering RNA (siRNA) against DCP2 or with a control siRNA (consists of a scrambled sequence that does not specifically target any transcript) 24 hours prior to transfecting the TTP and reporter plasmids. Quantitative reverse transcription PCR (RT-qPCR) assays showed that siRNA-mediated knockdown of DCP2 is effective and reduces DCP2 transcript levels to about 2% of its normal level (Figure S4A). Reduction of DCP2 levels further stabilizes the R-Luc-ARE reporter in HEK293 cells transfected with TTP_N-TZF_, but has no significant effect on the reporter levels in cells expressing full-length TTP (Figure 5B). We attribute this effect to the recruitment of DCP2 by the N-terminal activation domain of TTP, which in the absence of its C-terminal activation domain emerges as a more prominent means of mediating decay. While TTP_N_ mediates direct interactions with DCP2, it does not stimulate decapping activity *in vitro* (Figure S4B), suggesting that its predominant function is to facilitate the assembly of a decay-competent mRNP.

**Figure 5.**
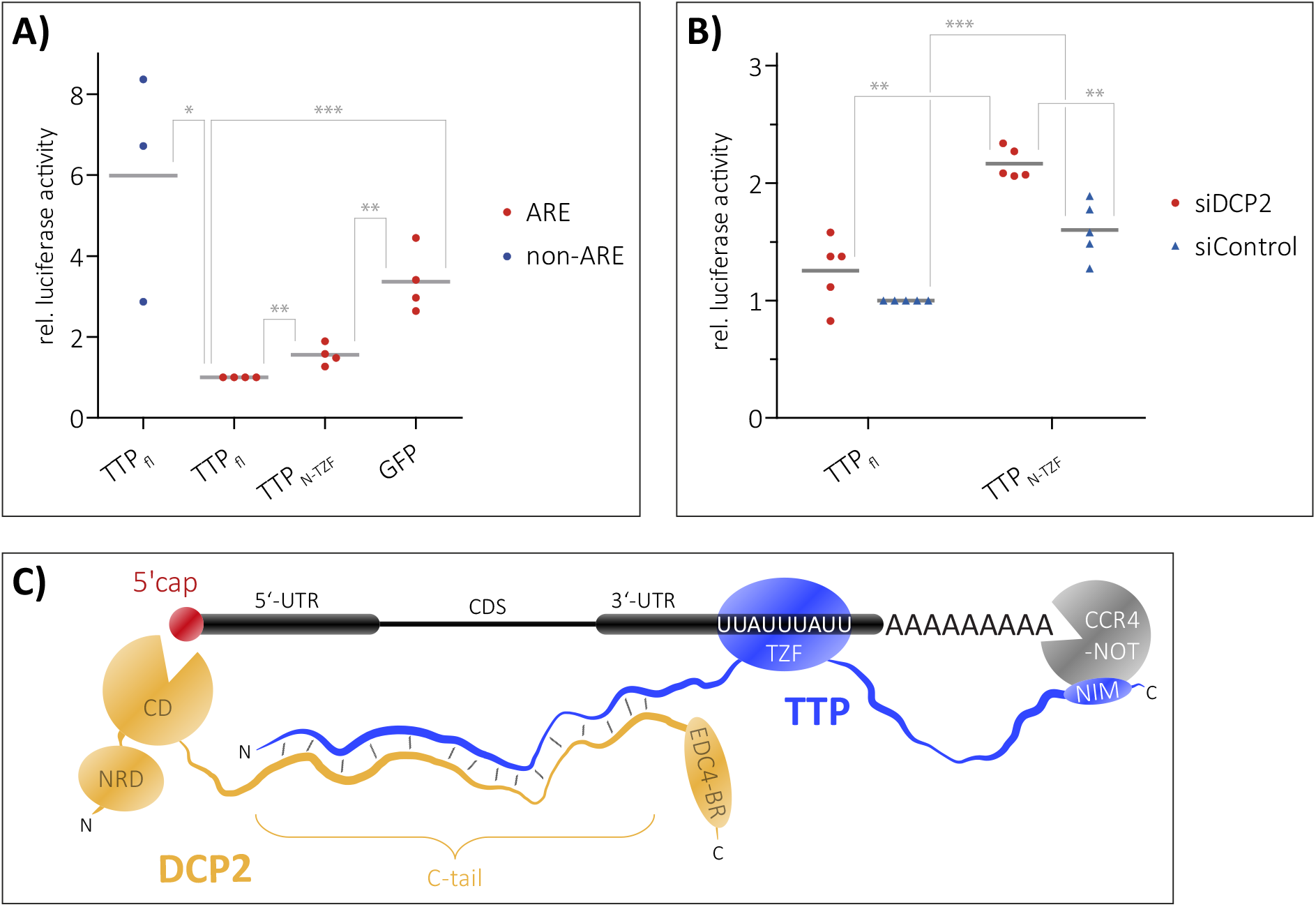
Recruitment of DCP2 by TTP_N_ contributes to decay of an ARE mRNA reporter. A-B) Luciferase reporter assays to determine the stability of an mRNA transcript containing AREs in the 3′-UTR. HEK293 cells were transfected with RLuc-ARE-A_90_-MALAT1 (ARE) reporter and plasmids expressing full-length TTP or TTP_N-TZF_. An FLuc-GFP reporter was used as a transfection control. An RLuc-A_90_-MALAT1 lacking the AREs (non-ARE) and GFP were used as negative controls in A. siDCP2 refers to siRNA-mediated knockdown of DCP2, while the siControl indicates treatment with a scrambled siRNA sequence that does not target any specific transcript in cells (B). Grey bars represent the mean of n independent experiments (n=4 for ARE reporter samples and n=3 for non-ARE reporter in A, n=5 in B). Individual data points are shown in all cases. The RLuc activity is normalized to FLuc; all RLuc/FLuc activities are expressed relative to that observed for full-length TTP (siControl treated sample for B), which was set to 1. Asterisks indicate the significance of the observed difference in each case as determined from p-values derived from unpaired t-tests. * p < 0.05 (significant), ** p< 0.01 (very significant), *** p<0.001 (extremely significant). A) The N-terminal activation domain of TTP is capable of mediating decay of the ARE reporter. A reporter lacking the ARE sequence (non-ARE) is not degraded by full-length TTP, and GFP does not degrade the ARE reporter, reaffirming that the observed degradation of the ARE reporter by the TTP proteins is a specific effect. B) Reduction of cellular DCP2 levels impedes the ability of TTP_N-TZF_ to mediate decay of the ARE reporter but does not significantly affect decay by full-length TTP. C) A model representing the modular interactions mediated by TTP that lead to the assembly of a decay-competent mRNP on an ARE mRNA. The TZF domain of TTP binds RNA, while the N- and C-terminal activation domains engage DCP2 (and possibly the decapping machinery) and the CCR4-NOT deadenylation complex, respectively. Bridging of events at the 5′-end and 3′-end by additional protein factors not shown in the figure (such as the LSm complex and PAT1) might further stabilize the weak TTP-DCP2 interactions.

## Discussion

The function of TTP in mediating rapid mRNA turnover hinges on its ability to recruit components of the mRNA degradation machinery to the target mRNA transcript. This is enabled by the modular domain organisation of TTP, comprising its central RNA-binding TZF domain and the flanking N- and C-terminal activation domains. Although the interactions of the TZF domain with an ARE-RNA as well as that of the C-terminal activation domain with the CCR4-NOT complex are fairly well characterized, the interactions mediated by the N-terminal activation domain have not been studied in as much molecular detail. Here we investigate the interaction of the TTP N-terminal activation domain (TTP_N_) with the decapping enzyme DCP2 *in vitro*. We show, using a combination of biochemical and biophysical methods, that TTP_N_ mediates a direct, but weak interaction with DCP2. This interaction is an unusual one as it involves IDRs of both interacting partners. This is in contrast to the typical examples of proteinprotein interactions, where the binding interface involves folded domains on both interaction partners or where an IDR of one protein interacts with a structured domain of its binding partner to form a well-ordered interface. A recent report on the binding of the linker histone (H1) to the nuclear protein prothymosin α (ProTα) highlighted the interaction between disordered regions of the two proteins, where both proteins retain their structural disorder in the complex (Borgia, Borgia et al. 2018). The interaction between two IDRs represents a new mode of biomolecular binding that has largely remained unexplored *in vitro* due to the focus of structural biology methods on protein order and structurally defined binding sites. The interaction between TTP_N_ and DCP2_C-tail_ is a fairly weak one, of a modest affinity of about 7 μM. Although the two proteins can be co-precipitated in GST-pulldown assays, they do not form a stable complex in size-exclusion chromatography, indicating rapid dissociation of the interacting partners from each other. Studies of protein-protein interactions, especially from a structural biochemistry perspective, have typically been dominated by strong, stoichiometric interactions that result in the formation of stable complexes. However, a high throughput study of the human interactome revealed that weak, sub-stoichiometric interactions in fact dominate the interactome and drive functionality in cells (Hein, Hubner et al. 2015). While strong interactions define core complexes that exist in isolation, weak interactions *in vitro* translate to transient, dynamic interactions in cells that can be regulated and remodelled easily. Assembly of decay-competent mRNPs on mRNA transcripts destined for targeted decay entail multiple weak interactions among components of the complex, which are then readily disassembled after decay of the mRNA (Gehring, Wahle et al. 2017). Using ARE-reporters, we show that TTP_N-TZF_, which lacks the C-terminal NIM, is still capable of mediating decay of an ARE-reporter, though not as efficiently as the full-length protein. A reduction of cellular DCP2 levels by siRNA knockdown does not affect full-length TTP but significantly hampers the ability of TTP_N_ to mediate decay. These observations suggest that the weak interaction between TTP_N_ and DCP2_C-tail_ is sufficient to induce degradation of a target transcript even in absence of the stronger interaction of TTP-NIM with the CCR4-NOT deadenylation complex. The interaction with DCP2 also incorporates an element of redundancy into TTP-mediated ARE-mRNA decay, which stringently regulates the levels of several transcripts involved in innate immunity (such as TNFα) (Khabar 2005). Interestingly, the eIF4E-transporter protein 4E-T, which was shown to interact with TTP in a proximity-dependent biotin identification (BioID) assay, bridges the 3′-end degradation machinery with the decapping machinery at the 5′-end, suggesting that TTP can assemble a decay-competent mRNP on its target transcript in multiple ways (Nishimura, Padamsi et al. 2015).

Weak multivalent interactions between proteins and/or RNA often lead to liquid-liquid phase separation *in vitro* and partitioning of proteins into membraneless organelles in cells. An example of such organelles in cells are cytoplasmic processing bodies (P-bodies) which are rich in mRNA decay factors (such as DCP1/2, EDC4, 4E-T and DDX6), but are now increasingly believed to be sites of mRNA storage for translationally repressed mRNA, rather than sites of decay (Aizer, Kalo et al. 2014, Kamenska, Simpson et al. 2016, Hubstenberger, Courel et al. 2017). The N- and C-terminal activation domains of TTP were reported to localize to P-bodies and recruit their mRNA cargo to these compartments (Franks and Lykke-Andersen 2007). We show that TTP_N_ does not undergo phase separation on its own at high concentrations, even in the presence of a molecular crowding reagent. However, DCP2 which is a core component of P-bodies has a strong tendency to undergo LLPS *in vitro*. The DCP2_ΔC_ construct encompassing the catalytic core and C-tail forms LLPS droplets at relatively low protein concentrations. Both the core and the unstructured C-tail of DCP2 also have a tendency to partition into droplets, albeit at higher concentrations than the DCP2_ΔC_ protein. The tendency of DCP2 to form LLPS droplets might stem from intra-molecular interactions of the DCP2 C-tail with itself or the catalytic core (Figures S3B and S2B, respectively). In the presence of equimolar amounts of DCP2_ΔC_ and DCP2_C-tail_, TTP_N_ also undergoes LLPS *in vitro*, an effect not observed with DCP2_core_, which lacks the TTP-binding region. These observations suggest that the weak interaction with DCP2 induces phase separation of TTP. This mechanism might lead to recruitment of TTP to P-bodies in cells, although it should be noted that TTP also interacts with other core components of P-bodies (Hubstenberger, Courel et al. 2017). The sequestering of TTP and DCP2 in phase-separated droplets *in vitro* or in P-bodies in cells also serves to increase their local concentration, which in turn would augment binding of the two proteins.

In summary, we show TTP uses its intrinsically disordered N-terminal activation domain to engage the unstructured C-terminal region of the decapping enzyme DCP2 in a weak, fuzzy-type interaction to mediate decay of its target ARE-mRNA (Figure 5C). This interaction is probably one of the many interactions mediated by TTP to assemble a transient decay-competent mRNP on the target transcript, and likely represents a common event in other targeted mRNA decay pathways. A very recent study by Tibble and co-authors show that decapping can be both activated and repressed in membraneless compartments, depending on protein-protein interactions that drive phase separation and induce conformational changes of *Schizosaccharomyces pombe* Dcp1/Dcp2 (Tibble, Depaix et al. 2021). Although TTP and DCP2 can partition in phase-separated droplets and have been found to be present in P-bodies in cells, it remains unclear as to where decay of the target mRNA eventually takes place and what releases TTP from P-bodies in cells. Our studies highlight the importance of investigating weak interactions, particularly between intrinsically disordered proteins (IDPs), to understand the dynamic assembly and disassembly of macromolecular complexes in mediating cellular functions.

## Materials and Methods

### Protein expression and purification

All protein sequences used in this study are of human origin.

#### TTP_N_

Residues 1-99 of TTP, corresponding to TTP_N_, were cloned into a modified pET28a vector bearing an N-terminal 6XHis-MBP tag, followed by a TEV protease cleavage site. The protein was expressed in *E. coli* BL21(DE3) gold pLysS cells grown in Terrific broth (TB). Protein expression was induced with 0.1 mM Isopropyl β-d-1-thiogalactopyranoside (IPTG) at 18°C. The culture was harvested 20 h post induction and the pellet was resuspended in lysis buffer (50 mM Tris pH 7.5, 1000 mM NaCl, 10 mM imidazole, 10% glycerol, 5 mM β-Mercaptoethanol) supplemented with 1 mM phenylmethylsulfonyl fluoride (PMSF) and 0.5 mg DNase I. Cells were lysed by sonication, clarified by centrifugation and filtration and subjected to Ni^2+^-affinity chromatography (IMAC) to isolate the target protein. Hìs-MBP-TTP_N_ was eluted from the Ni^2+^-affinity resin (Machery-Nagel #745400.100) in elution buffer (lysis buffer supplemented with 250 mM Imidazole). To cleave the His-MBP tag, TEV protease was added to the eluted protein to a final ratio of 1:50 (w/w). The protein mixture was dialysed to remove imidazole and subjected to a reverse Ni^2+^-affinity purification (the flow through containing primarily TTP_N_ and small amounts of His-MBP was collected). Following removal of His-MBP contaminants by incubation with Dextrin resin, the TTP_N_ protein was concentrated and loaded on a size-exclusion chromatography (SEC) column (HiLoad 26/60 Superdex 75 pg, GE Healthcare). Peak fractions from the first SEC step, performed in SEC buffer A (20 mM Tris pH 7.5, 1000 mM NaCl, 5% glycerol), were pooled and reloaded onto the SEC column to remove any traces of His-MBP contaminants. Peak fractions from the second SEC step, performed in SEC buffer B (50 mM Potassium phosphate pH 7.0, 50 mM NaCl) were pooled, concentrated and flash frozen in liquid nitrogen. In this and all other purifications, protein purity was monitored by SDS-PAGE analyses. The purification of TTP_N_ for FRET and LLPS is described in detail in Supplementary methods.

#### DCP2 proteins and GYF

DCP2_ΔC_ and DCP2_core_ were cloned into a modified pET28a vector with an N-terminal 6XHis-thioredoxin (Trx) tag, whereas DCP2_C-tail_ and GYF were cloned into pET28a with an N-terminal 6XHis tag. Proteins were expressed in *E. coli* BL21(DE3) Star pRare as described above for TTP_N_. All proteins were purified using a combination of Ni^2+^-affinity chromatography and SEC, using lysis buffer A and SEC buffer A. An intermediate heparin-affinity chromatography step (Hitrap Heparin column 5 mL) was additionally performed for DCP2_ΔC_. The affinity tags (6XHis or 6X-His-Trx) were removed by cleavage with HRV 3C protease prior to SEC.

#### GST-tagged proteins

GYF and DCP2 proteins were cloned into a modified pET vector bearing an N-terminal 6X-His-GST tag. The protease cleavage site between the GST-tag and the protein of interest was replaced by a 7-residue long glycine-serine-alanine rich linker, resulting in an uncleavable His-GST tag. The proteins were expressed and purified by Ni^2+^-affinity chromatography, as described above for Hìs-MBP-TTP_N_, followed by SEC (Hiload Superdex 200 16/600 for His-GST-DCP2_ΔC_/DCP2_core_ and Hiload Superdex 75 16/600 for His-GST-DCP2_C-tail_/GYF) in SEC buffer A. As with His-Trx-DCP2_ΔC_, an intermediate heparin-affinity chromatography step was performed for Hìs-GST-DCP2_ΔC_. Peak fractions were pooled and flash frozen as above. GST-SMG6_TPR_ (residues 520-1186) was a gift from Elena Conti (Max Planck Institute of Biochemistry, Martinsried).

### Fluorescent labelling of proteins

DCP2_C-tail_ was labelled with Cy3 (Cytiva Amersham Cy3 Maleimide Mono-Reactive Dye, product no.: PA23031) and TTP_N_ with Cy5 (Cytiva Amersham Cy5 Maleimide Mono-Reactive Dye, product no.: PA25031) for FRET experiments. Both constructs contained a single cysteine which was coupled to the dye via a maleimide reactive group. 200 nmol of each protein diluted in labelling buffer (20 mM HEPES pH 7.5, 100 mM NaCl, 100 μM TCEP) were pre-incubated at 37°C for 30 min before adding one vial of fluorescent dye resuspended in 50 μL DMSO. The reaction (total volume of 2 mL) was incubated at 25°C for 2 h, followed by an overnight incubation at 4°C. The excess dye was separated from the labelled protein by SEC (Superdex 75 10/300 GL, GE Healthcare) performed in SEC buffer D (20 mM HEPES pH 7.5, 100 mM NaCl). The SEC run was monitored at 280 nm to detect the protein as well as at 550 nm (for Cy3) or 650 nm (for Cy5) to track the fluorescent label. Peak fractions containing the labelled protein were pooled and concentrated. DCP2 proteins for fluorescence microscopy studies were labelled with maleimide-reactive XFD488 dye (AAT Bioquest #1878, identical to Alexafluor488) using a similar protocol as for Cy3-labelling of the protein. All buffers used for XFD488 labelling of DCP2 contained 150 mM NaCl. 100 nmol fluorescent dye was added to 100 nmol (DCP2_C-tail_) or 50 nmol (DCP2_core_ and DCP2_ΔC_) of protein in a total volume of 1 mL. The SEC run was monitored at 280 nm and 499 nm.

### FRET measurements

The experimental setup and analysis of FRET were performed according to (Mattiroli, Gu et al. 2018). Briefly, DCP2_C-tail_ was used as the FRET donor and TTP_N_ as the FRET acceptor in our setup. DCP2 was maintained at a fixed final concentration of 5 μM, while TTP was titrated in increasing concentrations, from 0-50 μM. In addition to the FRET reaction containing both labelled proteins, a donor bleed-through control (Cy3-DCP2_C-tail_ and unlabelled TTP_N_) and an acceptor direct excitation control (unlabelled DCP2_C-tail_ and Cy5-TTP_N_) were set up for each titration point. Proteins were mixed in a total volume of 35 μL in SEC buffer C (20 mM HEPES pH 7.5, 150 mM NaCl). Following overnight incubation at 4°C, samples were transferred to a black PerkinElmer OptiPlate 384-F and measured with a Tecon Spark plate-reader at 20°C (20 flashes, band width of 10 nm) in three different settings: FRET excitation at 520 nm and detection at 667 nm, donor bleed-through excitation at 520 nm and detection at 565 nm, acceptor direct excitation at 640 nm and detection at 667 nm. FRET values were corrected for donor bleed-through and acceptor direct excitation, after being adjusted against the buffer background. Values within a replicate series were normalized against the 35 μM TTP data point and fitted to a one site-specific binding model with Hill slope using GraphPad Prism 5.

### GST pulldowns

5-8 μg of GST-tagged bait proteins and 25 μg prey protein were mixed in GST-pulldown buffer (20 mM HEPES pH 7.5, 100 mM NaCl, 10% Glycerol, 0.1% NP40) to a total volume of 40 μL. Approximately 15% of the reaction was used as an input for the SDS-PAGE analysis. The reaction mixture was incubated at 4°C for 16 h, following which 12 μL of a 50% slurry of Glutathione Sepharose resin (GE Healthcare) was added. The mixture was further supplemented with 200 μL of GST-pulldown buffer and incubated at 4°C for 1 h. The beads were extensively washed with GST-pulldown buffer. Bound proteins were eluted in GST-pulldown buffer supplemented with 30 mM reduced glutathione and analyzed by SDS-PAGE and Coomassie or silver staining.

### Ni^2+^-affinity pulldowns

Fusions of TTP and DCP2 connected via a TEV protease cleavable linker were cloned into a modified pET28a vector bearing an N-terminal 6XHis-Thioredoxin tag. *E. coli* BL21(DE3) gold pLysS cells transformed with these plasmids were grown in 40 mL TB cultures at 37°C to an optical density at 600nm (OD_600nm_) of 2.5. Protein expression was induced with 0.5 mM IPTG at 18°C and cultures were harvested 20 h post induction. Pellets derived from a culture volume equivalent to a 1 mL culture at an OD_600nm_ of 20 were resuspended in 1.4 mL lysis buffer (20 mM Tris pH 7.5, 150 mM NaCl, 10 mM Imidazole) and lysed by sonication. Lysates were cleared by centrifugation and supernatants were mixed with 40 μL of equilibrated Ni^2+^-bead slurry (Machery-Nagel #745400.100) and incubated for 1 h at 4°C. Beads were washed three times with lysis buffer prior to addition of 40 μL of elution buffer (50 mM Tris pH 7.5, 150 mM NaCl, 500 mM Imidazole). Samples were incubated with elution buffer for 15 min at 30°C. Samples of total lysate, cleared (soluble) lysate and elution were analysed by SDS-PAGE.

### Analytical size-exclusion chromatography

All samples were prepared in a total volume of 40 μL in SEC buffer A supplemented with 2 mM DTT and analysed on a Superdex 75 3.2/300 Increase column (GE Healthcare). For the TTP-DCP2 interaction, TTP_N_ and DCP2_C-tail_ were mixed in equimolar ratios (ranging from 25-100 μM) and incubated overnight at 4°C. The individual proteins were also analysed by SEC.

The TTP-GYF interaction was analysed by mixing together 6.3 nmol TTP_N_ and 7.5 nmol GYF (1:1.2 molar ratio), followed by a 2 h-incubation at 4°C prior to SEC (Superdex 75 3.2/300 Increase column). As above, single proteins were also analysed by SEC (6.3 nmol each protein). Furthermore, the TTP_N_-GYF fusion construct, both in its single polypeptide form (intact fusion) or protease-cleaved form (resulting in two polypeptides), was analysed by SEC. A total of 80 μg of protein was analysed in each run.

### Luciferase and RT-qPCR assays

0.3×10^6^ HEK293 cells in DMEM media (Gibco #31966-021) supplemented with 10% fetal bovine serum were seeded in each well of 12-well plates. Replicates were set up for protein and RNA extraction for luciferase assays and RT-qPCR to determine the siRNA knockdown efficiency of DCP2, respectively.

Cells were transfected with 20 pmol siRNA against DCP2 (or a control siRNA, siScrambled) using ROTIFect (Carl Roth #P001.3), according to manufacturer’s instructions. 24 h post siRNA transfection, cells were transfected with 100 ng of either TTP or GFP expression plasmids and 500 ng of luciferase plasmids (Renilla-luciferase reporters containing or lacking the TNFα-ARE in the 3′-UTR, and Firefly-luciferase as a control) using PEI (Polysciences #24765). Cells were harvested 24 h post plasmid transfection in 200 μL lysis buffer (60 mM Tris pH 7.5, 30 mM NaCl, 1 mM EDTA, 1% Triton X-100) for protein extraction or 500 μL Trizol (Bio&Sell) for RNA extraction.

Luciferase assays were performed in duplicate for all biological replicates. Renilla and firefly luciferase assays were performed using commercially available reagents (Renilla-Juice Luciferase Assay #102531 and Beetle-Juice Luciferase assay Firefly #102511 from PJK GmbH) and activities were measured in a plate reader (Tecan GENios).

200 ng of DNA-free total RNA extracted from replicates was used for gene-specific cDNA synthesis by the MLV-reverse transcriptase (Qiagen). Quantitative (q)-PCRs were performed using the PowerUp SYBR Green Master Mix (ThermoFisher Scientific) on a Stratagene Mx3005P instrument. The data represent the mean of duplicates of one representative analysis. Primers used for reverse transcription and qPCR are listed below:

DCP2 fwd.: GCAGCAGAATTCTTTGATGAAGTG
DCP2 rev.: GCTGTCCCTCAGCATGTTCT (used for reverse transcription of DCP2 cDNA)
GAPDH fwd.: CTTCGCTCTCTGCTCCTCCTGTTCG
GAPDH rev.: ACCAGGCGCCCAATACGACCAAAT (used for reverse transcription of GAPDH)

### Liquid-liquid phase separation microscopy

Microscopy was performed with unlabelled proteins for phase contrast microscopy and Cy5-TTP_N_ or XFD488-DCP2 proteins for fluorescent microscopy. Individual proteins or protein mixtures were diluted to 2X the indicated concentration, incubated overnight at 4°C and were mixed with a crowding solution (30% PEG 8000, 100 mM Sodium citrate pH 7.0) in a 1:1 ratio immediately before the microscopy analysis.

#### Phase contrast microscopy

TTP_N_ and DCP2 constructs (DCP2_ΔC_, DCP2_core_ and DCP2_C-tail_) were diluted in SEC buffer C (20 mM HEPES pH 7.5, 150 mM NaCl) individually in different concentrations ranging from 495 μM to 20 μM or in equimolar mixtures of TTP and DCP2 (concentrations of 250, 100, 50 and 20 μM of each protein). A Zeiss Primo Star microscope with a Plan-Achromat 40x/0,65 Ph2 lens (with Ph2 condenser for phase contrast) was used to visualize formation of phase-separated droplets.

#### Fluorescence microscopy

Fluorescence microscopy was performed as above, using fluorescently-labelled TTP_N_ and DCP2 proteins. Proteins were diluted and mixed together in a 1:1 ratio with 180 μM of each protein and incubated overnight at 4°C. A Zeiss Axio Observer 7 ApoTome microscope with a Plan-Apochromat 63x/1.4 DIC oil immersion lens was used to visualize droplets containing the two proteins. DCP2 was detected via the X-channel (475 nm filter) and TTP was detected via the Y-channel (630 nm filter). Optical sectioning was performed using the Zeiss Apotome to improve the contrast of the fluorescent droplets by eliminating out-of-focus light and reducing the background from multiple focal planes.

## Supplemental Information

Supplemental information (methods and figures) is available for this manuscript.

## Acknowledgements

We thank Jens Lykke-Andersen for the gift of the human DCP2 plasmid and Christian Freund for the human GIGYF2 plasmid. The luciferase reporter plasmids were a generous gift from Catia Igreja and the former Izaurralde group. We would like to acknowledge the assistance of the Core Facility BioSupraMol (Freie Universität Berline) supported by the DFG. We thank Tanja Matković-Rachid and Francesca Bottanelli for sharing their expertise on optical microscopy experiments, and Eric Simko and Eugene Valkov for the *in vitro* decapping assay. We are grateful to Janosch Henning and Cecilia Perez-Borrajero as well as members of our laboratory for helpful discussions and comments on the manuscript. This study was supported by the Priority Programme SPP 1935 of Deutsche Forschungsgemeinschaft (CH1245/3-1 and CH1245/3-2 to S.C.). The authors declare that there are no conflicts of interest.

## Author contributions

V.D.M. purified the proteins used in this study with assistance from N.M. and S.C. Biochemical assays and microscopy experiments were done by V.D.M. FRET assays and SEC-MALS were performed by N.M. and V.D.M. Luciferase assays were carried out by T.D., S.C. and V.D.M. S.C. and V.D.M designed the study, and wrote the manuscript with input from N.M. and T.D.

