## Supplemental information for "Intrinsically disordered regions of Tristetraprolin and DCP2 directly interact to mediate decay of ARE-mRNA"

#### Supplemental Methods

*Protein purification of $TTP_N$ for FRET and LLPS* microscopy

We exploited the lack of interaction between $TTP_N$ and $DCP2_{\Delta C}$ construct, despite the stabilizing effect of DCP2 on TTP, to purify pure homogenous $TTP_N$ for fluorophore labelling (Figure S2C, flow chart). To this end, we used a fusion construct of $TTP_N$ and $DCP2_{\Delta C}$, TTP(1-99)-3C-DCP2(1-353), cloned into a modified pET28a vector with an N-terminal 6X-His tag. The protein was expressed in *E. coli*, resuspended in lysis buffer B (50 mM Tris pH 8.0, 1000 mM NaCl, 10 mM imidazole, 10% glycerol, 5 mM β-Mercaptoethanol) and subjected to $Ni^{2+}$-affinity chromatography as described in the method section of the main text (Figure S2C, lane 1). The protein obtained from $Ni^{2+}$-affinity chromatography was dialysed overnight to reduce the concentration of NaCl and imidazole (Figure S2C, lane 2) and loaded onto a heparin column (HiPrep Heparin FF 16/10, GE Healthcare). The protein was eluted off the column using a salt gradient ranging from 100-600 mM NaCl (Figure S2C, lane 4). Peak fractions were pooled and treated with HRV 3C protease at 4°C, while undergoing dialysis simultaneously to lower the concentration of NaCl in the protein mixture. Treatment with HRV 3C protease resulted in cleavage of the TTP-DCP2 fusion as well as the His-tag (Figure S2C, lane 5). To separate the two cleaved polypeptides, the dialysed sample was once again loaded on a heparin column (HiTrap Heparin HP 5 mL, GE Healthcare). $TTP_N$ flows through the column as it has no affinity for the heparin resin (Figure S2C, lane 6) and is separated from the heparin column-bound DCP2 (Figure S2C, lane 7). The flow through of the heparin column was concentrated and further purified by SEC (HiLoad 26/60 Superdex 75 pg, GE Healthcare) in SEC buffer containing 20 mM Tris pH 7.5, 150 mM NaCl. The peak fractions from SEC were pooled, concentrated and flash frozen in liquid nitrogen (Figure S2C, lane 8).

*Decapping assay*

Full-length DCP2 with an N-terminal MBP fusion and C-terminal His-tag was produced using the pnEA-NpM backbone and expressed in *E. coli* BL21 Star (DE3). Following lysis in 50 mM HEPES pH 7.0, 300 mM NaCl, 5% glycerol, protein was purified by affinity purification using amylose resin followed by immobilized metal affinity chromatography and polished by gel filtration with a HiLoad 26/600 Superdex 200 pg column.

*In vitro* synthesized RNA (127 nucleotides) was labelled with [γ-32P]GTP using Vaccinia Capping Enzyme (NEB) and 2'-O-Methyltransferase (NEB) according to manufacturer's protocol. Labelling reactions contained 1 µg of RNA and 5 µL of 3000 Ci/mmol [γ-32P]GTP. Unincorporated GTP was removed using a G-50 spin column and capped RNA was purified by phenol:chloroform extraction followed by LiCl precipitation. Labelled RNA was resuspended in 100 µL water.

Decapping reactions were carried out in 50 mM Tris pH 7.5, 50 mM ammonium sulphate, 0.1% BSA, 5 mM $MgCl_2$, and 5 mM $MnCl_2$. 10 µL cap-labelled RNA was used in 50 µL decapping reactions with 200 nM MBP-tagged full-length DCP2 and $TTP_N$ was added at indicated concentrations. Reactions were carried out at 10°C for 1 hour and timepoints were taken in 10 min increments and quenched with EDTA. TLC plates (PEI Cellulose F, 20 cm x 20 cm, Millipore), pre-run in water and scored to produce 1 cm lanes, were spotted with 1 µL of timepoint samples and developed with 0.75 M LiCl. BAS-IP phosphor screens (Fuji) were exposed to TLC plates overnight and imaged with an Amersham Typhoon (GE) and signal was quantified using ImageQuant (Cytiva).

### Supplemental Figures

#### Figure S1 (related to Figure 1 of the main text)

A)

```
TTP         ------------------------------------------------------------
BRF1     1 MGRARKSGLGQRSRPTASRSEAAVQPGVRKARGAGNWRVGLQTGEAAPSPHRDLRDTPDP   60
BRF2        ------------------------------------------------------------

TTP      1 --------------MDLTAIYES---------LLSLSPD--------VPVPSDHGG---T   26
BRF1    61 RPWLARTHRMTTTLVSATIFDLSEVLCKGNKMLNYSAPSAGGCLLDRKAVGTPAGG----  116
BRF2     1 ---------MSTTLLS--AFYDVDFLCKTEKSLANL---NLNNMLDKKAVGTPVAAAPSS   46
                          .   :            *                * :  ..

TTP     27 ESSPGWGS----SGPWSLSPSDSSPSGVTSRLPGRSTSLVEGRSCGWVP-----------   71
BRF1   117 ----GFPRRHSVTLPSS-------------KF---------------------HQNQLLS  139
BRF2    47 GFAPGFLRRHSASNLHALAHPAPSPGSCSPKFPGAA----NGSSCGSAAAGGPTSYGTLK  102
               *:      :   :             ::

TTP     72 PPP-----------------GFAPLAPRLG--PELSPSPTSPTATSTTPSRYKTELCRTF  112
BRF1   118 SLKGEPAPALSSRDSRFRDRSFSEGGERLL------PTQKQPGGGQVNSSRYKTELCRPF  192
BRF2   103 EPSGGGGTALLNKENKFRDRSFSENGDRSQHLLHLQQQQKGGGGSQINSTRYKTELCRPF  162
                               .*:  . *           .   . . . :******** *
                               TZF
TTP    113 SESGRCRYGAKCQFAHGLGELRQANRHPKYKTELCHKFYLQGRCPYGSRCHFIHNPSEDL  172
BRF1   193 EENGACKYGDKCQFAHGIHELRSLTRHPKYKTELCRTFHTIGFCPYGPRCHFIHNAEERR  252
BRF2   163 EESGTCKYGEKCQFAHGFHELRSLTRHPKYKTELCRTFHTIGFCPYGPRCHFIHNADERR  222
           .*.* *:** *******: ***. .**********:.*:  * **** ******* .*

TTP    173 AAPG-----------------------HPPVLRQSISFSGLPSGRRTSPPPPGLAGPSLS  209
BRF1   253 ALAGARD-----------------LSADRPRLQHSFSFAGFPSAAATA------AATGLL  289
BRF2   223 PAPSGGASGDLRAFGTRDALHLGFPREPRPKLHHSLSFSGFPSGHHQP--PGGLESPLLL  280
              .                         * *::*:**:*:**.           .  *

TTP    210 SSSFSPSSSPPPPGDLP---LSPSAFSAAPGTPLARRDPTPVCCPSCRRATPISVWGPLG  266
BRF1   290 DSPTSI--TPPPILSADDLL------------------GSPT------------------  311
BRF2   281 DSPTSR--TPPPPSCSSASSCSSSASSCSSASAASTPSGAPTCCASAAAAAAAALLYGTG  338
           .*  *   :***                           :*.

TTP    267 G---------------------------------LVR-----------------------  270
BRF1   312 ----------------LPDGTNNPFAFSSQELASLF------------------------  331
BRF2   339 GAEDLLAPGAPCAACSSASCANNAFAFGPE-LSSLITPLAIQTHNFAAVAAAAYYRSQQQ  397
                                             *.

TTP    271 ------------------------------------------------TPSVQSLGSDPD  282
BRF1   332 --------------APSMGLPGGG-----SPTTFLFRPMSESPHMFDSPPSPQDSLSDQE  372
BRF2   398 QQQQGLAPPAQPPAPPSATLPAGAAAPPSPPFSFQLPRRLSDSPVFDAPPSPPDSLSDRD  457
                                                            **  .  ** :
                                                   NIM
TTP    283 EYASS---GSSLGGSDSPVFEAGVFAPPQPVAAPRRLPIFNRISVSE- 326
BRF1   373 GYLSSSS--SSHSGSDSPT-----------LDNSRRLPIFSRLSISDD 407
BRF2   458 SYLSGSLSSGSLSGSESPS-----------LDPGRRLPIFSRLSISDD 494
            * *.    .* .**:**            :   ******.*:*:*:
```

B)

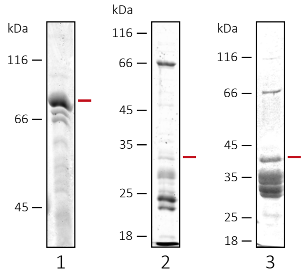


1) His-Trx-TTP$_{fl}$-GFP

2) His-Trx-TTP$_{N}$-Sumo

3) His-GST-TTP$_{N}$-His

A) Multiple sequence alignment of human TTP (TIS11), BRF1 (ZFP36L1/TIS11a) and BRF2 (ZFP36L2/TIS11d). Invariant residues are coloured dark red and highly conserved residues are in light red. The degree of conservation is shown by the symbols *, : and . which denote invariant, highly conserved and weakly conserved, respectively. The tandem zinc finger (TZF) domain and the Not1-interacting motif (NIM) are well-conserved across the paralogues while the N- and C-terminal activation domains diverge considerably in length and sequence. B) SDS-PAGE analyses of purification attempts of full-length TTP (left panel) and the TTP N-terminal activation domain ($TTP_N$, middle and right panels), expressed as N- and C-terminal fusions with diverse solubility and stability tags such as Sumo, Thioredoxin (Trx), GFP and GST. The approximate position of the intact protein on the gel is indicated by a red bar. TTP is heavily degraded upon cell lysis, even when fused to large tags.

**Figure S2 (related to Figure 2 of the main text)**

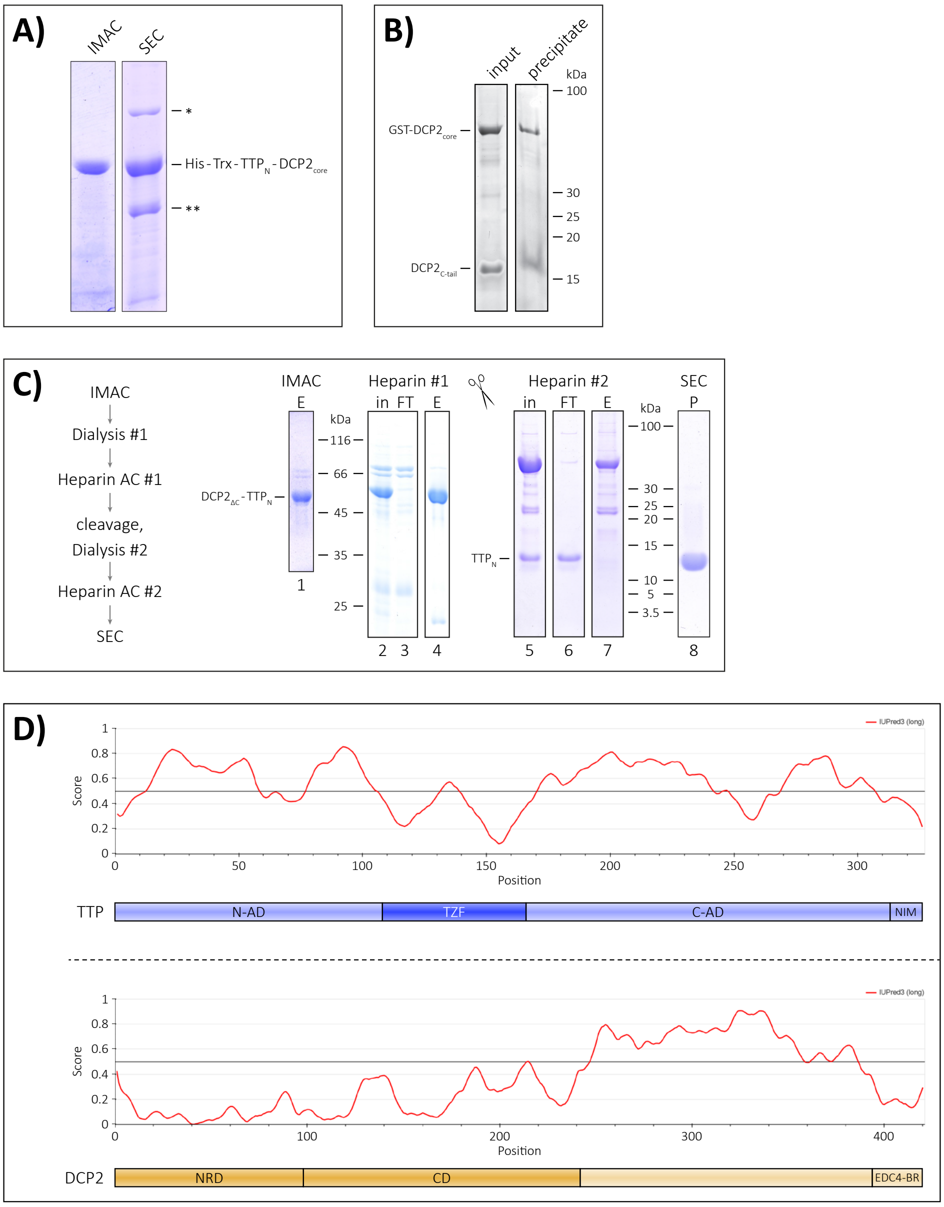


A) SDS-PAGE analyses of large-scale immobilized metal ion affinity chromatography (IMAC, $Ni^{2+}$-affinity purification) and size-exclusion chromatography (SEC) of the $TTP_N$-$DCP2_{core}$ fusion protein. Although fusion of $TTP_N$ with the $DCP2_{core}$ yields apparently stable protein in a small-scale $Ni^{2+}$-affinity purification, the protein is prone to degradation and cannot be purified to homogeneity on a large scale. * indicates a chaperone contaminant and ** indicates degradation products of the $TTP_N$-$DCP2_{core}$ fusion protein. These observations suggest that fusion of the DCP2 core to $TTP_N$ does not rescue the proteolytic instability of TTP. B) GST-pulldown assays of GST-$DCP2_{core}$ to the $DCP2_{C\text{-}tail}$. The C-tail of DCP2 folds back upon the DCP2 core, precluding its interaction with $TTP_N$ in solution. The left and right panels denote input and precipitate, respectively. Proteins are visualized by silver staining. C) Schematic and SDS-PAGE analyses of purification of $TTP_N$ from the $TTP_N$-$DCP2_{\Delta C}$ fusion. In - input, FT – column flowthrough, E – elution, P – peak of SEC purification. The fusion protein is stable and is purified to homogeneity using a combination of IMAC ($Ni^{2+}$-affinity chromatography) and Heparin-affinity chromatography (Heparin AC #1). Cleavage of the fusion produces stable $TTP_N$ and $DCP2_{\Delta C}$ which are separated from each other by a second Heparin-affinity chromatography step (Heparin AC #2). The column flowthrough containing relatively pure $TTP_N$ is concentrated and subjected to SEC to obtain a homogenous preparation of the protein. D) Prediction of intrinsic disorder of TTP and DCP2 using the program IUPRED (https://iupred.elte.hu/). TTP is mostly intrinsically disordered, except for its central TZF domain. The NRD-CD catalytic core of DCP2 is followed by a long, disordered stretch of 170 residues. IDRs of both proteins contain known protein-protein interaction motifs.

**Figure S3 (related to Figure 3 of the main text)**

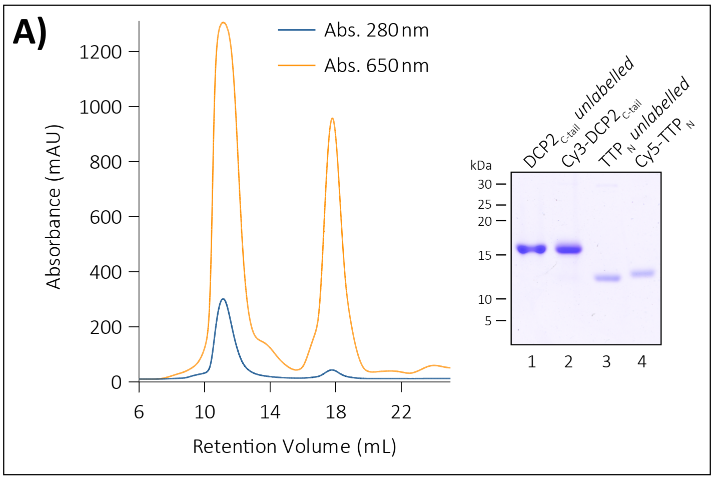


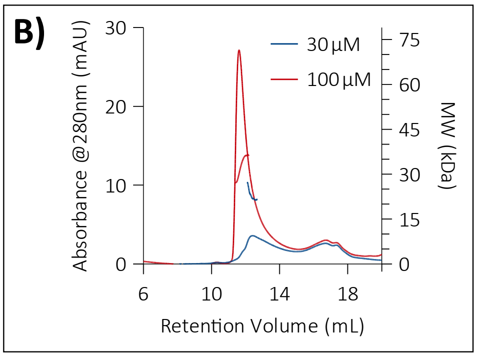


A) SEC purification of the products of the Cy5-labelling reaction of $TTP_N$ to separate the fluorescent dye from the labeled protein. The run is monitored at 280 nm and 650 nm to track the protein and the fluorescent label. A similar reaction and SEC purification were performed to obtain Cy3-labelled $DCP2_{C\text{-}tail}$. An SDS-PAGE analysis of the labelled and unlabeled TTP and DCP2 proteins used in the FRET assay is shown. B) Multi-angle light scattering in tandem with size-exclusion chromatography (SEC-MALS) analysis of DCP2 at 25 µM and 100 µM to determine its concentration-dependent oligomerization. DCP2 is a monomer at low concentration (25 µM) as evident from the experimental molecular weight (approximately 15 kDa) but displays a tendency to form higher order oligomers (30-40 kDa) at a high concentration (100 µM).

**Figure S4 (related to Figure 5 of the main text)**

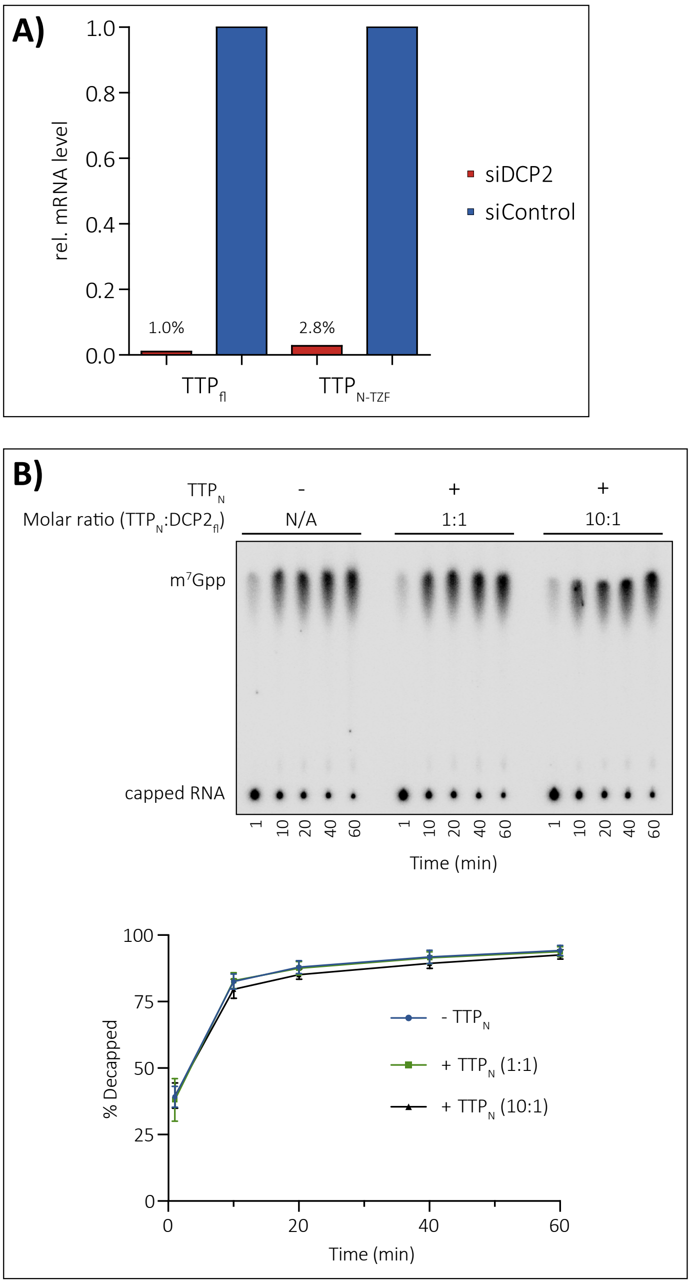


A) RT-qPCR assays to determine the level of DCP2 mRNA in cells treated with an siRNA against DCP2 (DCP2 k.d.) and a control siRNA (a scrambled sequence that does not target any transcript). The data shown here is a representative set out of 5 independent experiments where the relative luciferase activity of an ARE mRNA reporter was determined in the presence of different TTP constructs in cells containing or lacking DCP2 (Figure 5B). B) *In vitro* decapping activity of DCP2 in the presence of TTP$_N$. The direct effect of TTP$_N$ on DCP2 activity was tested *in vitro* with TTP added equimolar and in 10-fold excess to full-length DCP2. Decapping reactions were terminated at indicated time points and cleaved m$^7$GDP product was resolved from capped RNA substrate by thin layer chromatography (top panel). Signal was quantified and product was expressed relative to total as "% decapped" (bottom panel). Addition of TTP$_N$ does not affect the decapping activity of DCP2 *in vitro*.
